# ADRA2B genetic polymorphism modulates vagus nerve stimulation response

**DOI:** 10.64898/2026.11.03.763718

**Authors:** Elizabeth Riley, Senegal Mabry, Mary MacMillan, Lindsey Ko, Frank A. Middleton, Eve De Rosa, Adam Anderson

## Abstract

**Introduction:** Given interest in vagus nerve stimulation (VNS) as a potential therapy for Alzheimer’s disease, due to its hypothesized influence on the noradrenergic (NA) system, there is a need to understand the pronounced individual differences in responsiveness to VNS in aging. We tested whether a genetic polymorphism related to NA signaling, the Δ301-303 deletion in the ADRA2B receptor, modulates transcutaneous auricular (taVNS) effects on task-related pupillary responses or pupil-linked cortical activity in noradrenergic projection territory.

**Methods:** Forty-one adults (61% deletion carriers; ages 19-79) performed a visual oddball task during a sham-controlled single-blind crossover taVNS trial (NCT06880510). Of these people, 25 (68% deletion carriers) also performed the same task with simultaneous fMRI.

**Results:** ADRA2B genotype predicted the taVNS effect on oddball pupillary responses (t(37) = 3.03, p = .004, d = 0.84). Deletion carrier responses were more likely to increase, while non-carrier responses decreased. Carriers showed stronger spatial alignment between pupil-BOLD coupling and cortical norepinephrine transporter (NET) expression (t(22) = 2.82, p = .010, adjusted d = 1.21). This difference was selective to NET among 19 PET maps (all other p > .11). The individual effect of taVNS predicted NET alignment of pupil-BOLD coupling (ρ = .45, p = .023).

**Conclusions:** ADRA2B Δ301-303 deletion status, which is thought to increase NA availability, predicts the direction of the pupillary response to taVNS and how tightly pupil-linked BOLD activity aligns with cortical noradrenergic projection territory.

## Introduction

Pupil dynamics provide a noninvasive window into central nervous system arousal and have been linked to widespread cortical activity using functional magnetic resonance imaging (fMRI)(1–3). The neurochemical system most strongly associated with pupil dynamics is the locus coeruleus (LC) and noradrenaline (NA) system, but whether genetic variation in the LC-NA system affects pupillary responses or their cortical correlates is unknown. Understanding this may have clinical relevance, given increased interest in interventions such as vagus nerve stimulation (VNS), particularly transcutaneous auricular VNS (taVNS), that engage LC-NA pathways and are being considered a potential therapy for Alzheimer’s disease (AD) (4,5). This is largely because of the known anatomical connectivity between the vagus nerve, which carries critical information about the body’s internal physiological state, and LC, which receives direct input from the vagus nerve’s main central integration site, the nucleus of the solitary tract (6). The LC degenerates early in the course of AD (7–11), Parkinson’s disease (12–14), Progressive Supranuclear Palsy (15), and other neurodegenerative diseases (16,17). taVNS is hypothesized to support LC function by increasing NA release (4,18). This hypothesis is strongly supported by the known role of the LC/NE system in mediating the effects of VNS in other clinical contexts: epilepsy (19–23), migraine (24–29), and depression (30). However, efforts to directly evaluate the effects of taVNS on the LC have so far been hampered by inconsistent and contradictory results. Using fMRI, pupillometry, salivary alpha amylase, and EEG, VNS and taVNS has been shown to increase (31–34), decrease (29,35,36), alter in some other way (18,37–39), or not affect (40) LC activity or its physiological correlates. Data about these effects in older adults is sparse, but early reports suggest that the effect of taVNS may not be the same across age groups (36), with even more differences in those with neurodegenerative disease.

To resolve these conflicts, understand the mechanistic effect of vagus nerve stimulation on the LC-NA system, and potentially make effective use of taVNS in aging, the potential sources of variability in taVNS effects must be understood. Conversely, variability in taVNS responsiveness may also tell us something about the state of LC or brain activity and allow us to predict who may benefit from taVNS. One source of interindividual variability in LC-linked cognitive processes, as well as LC-linked cognitive aging outcomes, is a Δ301–303 polymorphism in the ADRA2B receptor protein, a metabotropic receptor that modulates noradrenergic signalling. The α2B-adrenergic receptor is expressed presynaptically in brain and vasculature and contributes to the modulation of adrenergic release via a negative feedback mechanism (41,42), which the Δ301–303 deletion variant (43) is thought to alter (44–46). Its behavioral correlates, including enhanced emotional memory, perception and attention (47–50) and altered responses to stress and noradrenergic drugs (51,52), together with supporting in vitro evidence (44), suggest it increases NA availability. It has also been shown, in a small sample, to have favorable effects on cognitive aging (53).

Here, we investigate the role of the ADRA2B polymorphism on taVNS responsiveness with sham and verum taVNS across an adult lifespan sample, a genotype-dependent partial reanalysis of our single-blind active sham-controlled crossover trial (NCT06880510) (36,54). To characterize whether individual differences in pupil-linked cortical activity reflect variability in NA signalling, we compared participant specific pupil-BOLD (blood oxygen level dependent) coupling maps and a PET-derived map of norepinephrine transporter (NET) expression. We then tested whether this spatial alignment differed by ADRA2B genotype. To our knowledge this is the first investigation of genetic predictors of noninvasive VNS responsiveness.

## Methods

### Participants

All participants were enrolled in a study on healthy brain aging at Cornell University. Forty-one participants participated in our taVNS clinical trial (54) and were genotyped. Of those, 25 also completed fMRI.

### Relationship to previously reported results

Of the 41 participants, results from 27 of them were reported previously (36) along with full CONSORT reporting. Fourteen participants were tested afterwards under the same protocol (mean age 55.0 y, SD = 13.2). Eleven participants from the earlier report are not included: 10 provided no saliva sample for genotyping, and 1 had insufficient usable oddball trials.

### Study procedures

Participants were screened for diagnosed cognitive impairment, neurological disease, severe head injury, ocular disease, cardiac arrhythmia, or any MRI contraindication, and had vision and hearing that were normal or correctable to normal. Participants were instructed to get a full night of sleep, abstain from alcohol the night before testing, and drink their habitual amount of caffeine, ending 1 h before testing. Testing consisted of taVNS, MRI, or both (see **Fig. 1, Table 1**), plus assessment of cognition with NIH Toolbox, saliva collection, and surveys including the demographics reported. The study was covered under Cornell Institutional Review Board protocol 1910009087.

**Table 1.**
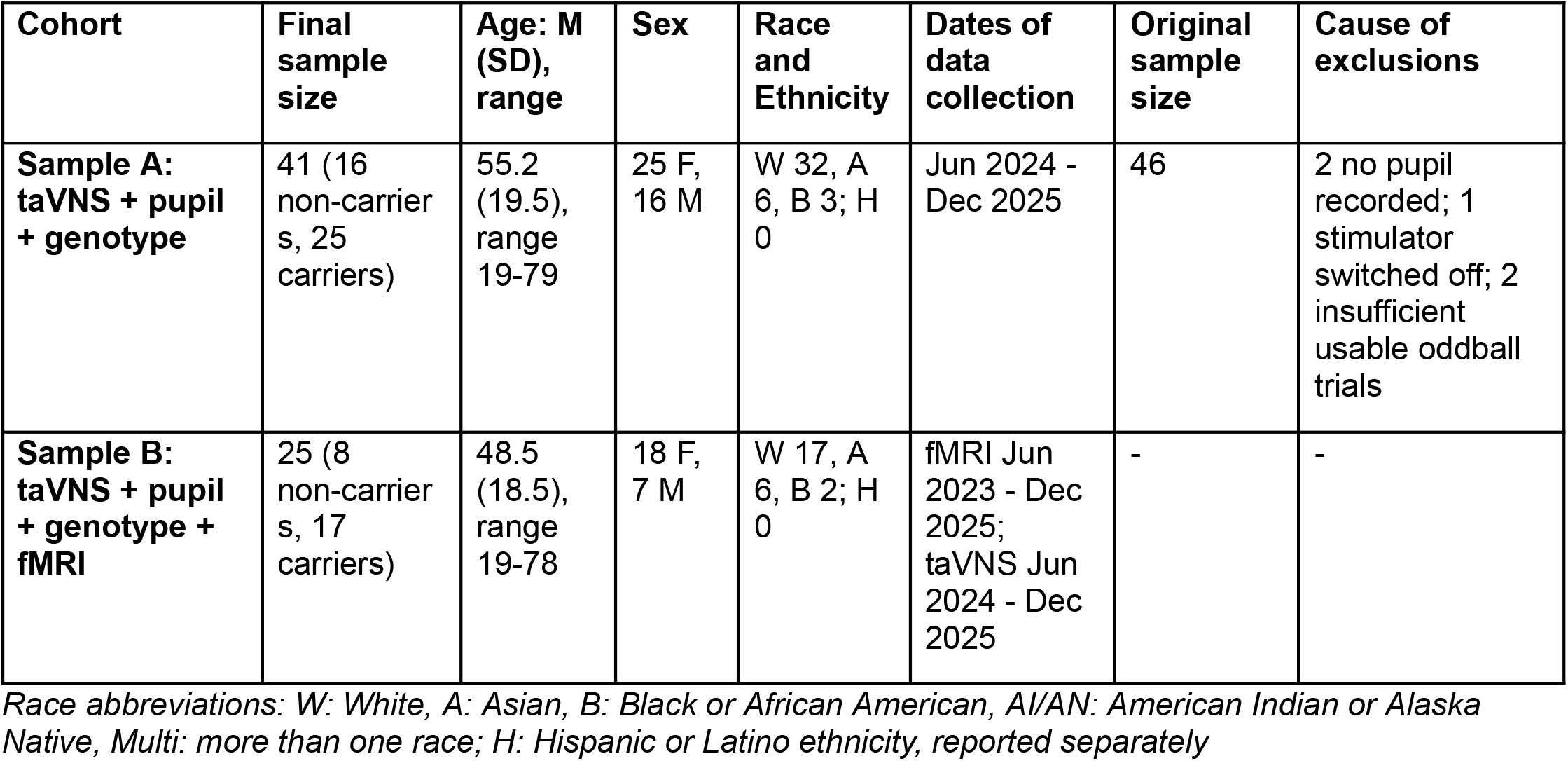
Participant information and demographics.

**Table 2.** taVNS effect on the oddball pupil response: specificity, robustness and group comparability.

| Category | Check | Model | Deletion carrier effect [95% CI] | p for deletion carrier effect |
|---|---|---|---|---|
| — | Primary result | Verum-Sham oddball AUC ~ carrier + age + order | b = 9.28 [3.08, 15.49] | **.004; adjusted Cohen's d = 0.84** |
| **Specificity** | Standard trials | Verum-Sham standard-trial AUC ~ carrier + age + order | b = 0.36 [−2.75, 3.47] | .82 |
|  | Bright trials | Verum-Sham bright-trial AUC ~ carrier + age + order | b = 0.68 [−4.21, 5.58] | .78 |
|  | Sham session only | Sham oddball AUC ~ carrier + age | b = −2.05 [−9.16, 5.05] | .56 |
|  | Verum session only | Verum oddball AUC ~ carrier + age | b = 7.97 [1.25, 14.70] | **.021** |
| **Robustness** | Strongest 1, 2, 3 carrier responses removed | Primary model | b = 8.29 / 7.41 / 6.51 | **.007 / .008 / .010** |
|  | Noisiest 1, 2, 3 participants removed | Primary model; noise = mean absolute 1-ms change in pupil diameter (mm), noisier session per person | b = 9.86 / 9.56 / 9.99 | **.003 / .004 / .003** |
|  | Leave-one-out | Primary model, each participant removed in turn (41 fits) | b = 8.29 to 10.11 | **.002 to .010 (41 of 41 < .05)** |
|  | Rank-based statistics | Mann-Whitney U = 104 | rank-biserial r = 0.48 | **.011** |
|  | Nonparametric statistics (permutation test) | Primary model, carrier labels shuffled 20,000 times | b = 9.28 | **.005** |
|  | Data preprocessing cutoff | Primary model, measures rebuilt interpolating over longer gaps (750 ms) | b = 8.73 [2.61, 14.86] | **.006** |
|  | All between-subjects covariates at once | Primary model + sex + race + current difference + washout; N = 34 (13 / 21) | b = 8.82 [0.82, 16.82] | **.032; adjusted Cohen's d = 0.79** |
|  | Sham oddball response size as covariate | Verum oddball AUC ~ carrier + sham oddball AUC + age + order | b = 8.57 [3.07, 14.07] | **.003** |
| **Group comparability** | Baseline pupil diameter | Sham-session median pupil diameter ~ carrier + age; N = 43 (17 / 26) | b = −0.03 mm [−0.37, 0.30] | .85 |
|  | Stimulation current, Sham | Welch t-test; N = 35 (14 / 21) | carriers − non-carriers = −305 µA | .51 |
|  | Stimulation current, Verum | Welch t-test; N = 36 (14 / 22) | −331 µA | .33 |
|  | Current difference (Verum-Sham) | Welch t-test; N = 35 (14 / 21) | −7 µA | .98 |
|  | Washout between sessions | Welch t-test; N = 35 (13 / 22) | +0.1 min | .98 |
|  | Treatment order | χ² test, carrier × session order (non-carriers 9 Sham-first / 7 Verum-first; carriers 12 / 13) | χ²(1) = 0.04 | .85 |

**Fig. 1.**
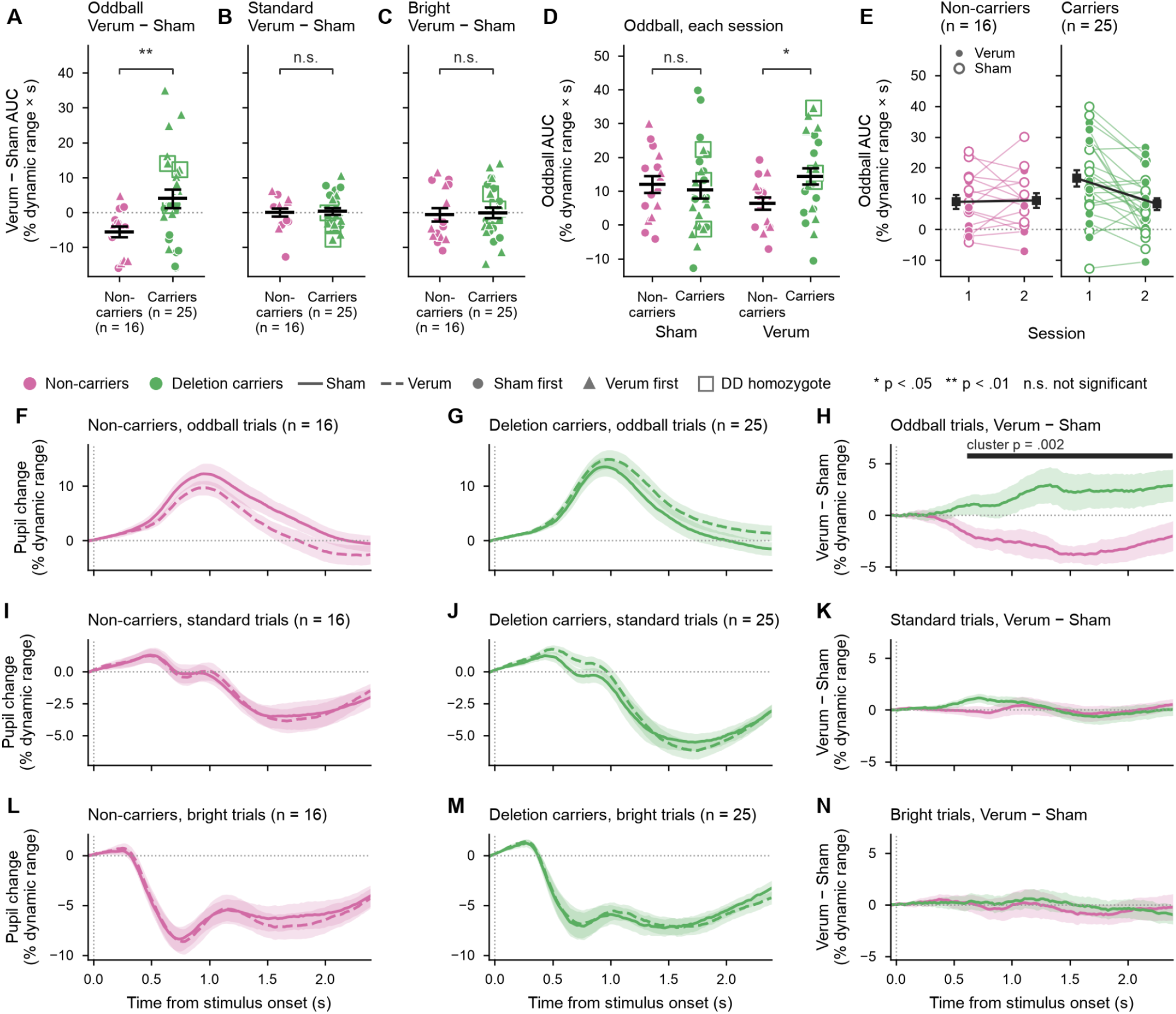
Effect of taVNS on task-evoked pupillary responses by ADRA2B carrier status (N = 41; 16 non-carriers, 25 carriers). (A-C) Verum - Sham AUC (% dynamic range × s) for oddball (A), standard (B) and bright (C) trials; brackets show significance of carrier effect from Verum − Sham AUC ∼ carrier + age + treatment order. (D) Oddball AUC in each session; brackets show significance of carrier effect from AUC ∼ carrier + age within each session. Mean ±1 SEM shown. (E) Oddball AUC in first and second sessions in non-carriers and carriers; closed vs. open circles show whether stimulation was Verum or Sham. (F-N) Mean pupil traces (% dynamic range; ribbons, ±1 SEM) for oddball (F-H), standard (I-K) and bright (L-N) trials: Sham and Verum in non-carriers (F, I, L) and carriers (G, J, M), and Verum - Sham by group (H, K, N); in H, the bar marks the significant cluster from a cluster-based permutation test of the carrier term (Verum - Sham ∼ carrier + age + treatment order at each 20 ms step, 5,000 permutations; corrected p shown). Marker shape shows which session came first; open squares mark DD homozygotes. ^*^ p < .05, ^**^ p < .01.

### Oddball task details

Participants completed a visual oddball task with pupillometry (36,55). The oddball task had three types of stimuli presented for 75 ms each: standard (60%, smaller purple circles isoluminant with the background), oddball (20%, larger purple circles isoluminant with the background, requiring a button press), and bright (20%, standard-size purple circles brighter than the background). Between stimuli, participants fixated on a black dot (0.15° visual angle). For the fMRI participants, the inter-stimulus interval was jittered (mean = 2.0 s, median = 1.5 s, range = 0.5-11.0 s, sequence designed with optseq2) and the total length was 10 minutes. For the taVNS participants, the inter-stimulus interval was jittered (2.5 - 3.5 s, uniform), the total length was 8 minutes, and the task was completed twice, as described below.

### taVNS administration

Each participant completed single-blind verum and sham sessions with transcutaneous electrical stimulation delivered with a tVNS Technologies research stimulation device. Stimulation parameters were: 25 Hz, 250 µs pulse width, 50% duty cycle (30 s on, 30 s off). Sham location: left earlobe; verum location: left cymba concha. The order of the sham and verum sessions was counterbalanced. The current level was adjusted individually for strong sensation (36). During the washout period (mean length 32.0 min, SD = 9.7), participants underwent cognitive testing.

### Integrity of the blind

The researchers who placed the tVNS Technologies device were aware of which position was verum and which was sham. The participants were not aware that either position was a sham. Pupillary and fMRI data was analyzed before receiving genotype data.

### Pupillometry

Eye movements and pupil size were recorded with an Eyelink 1000 Plus eye tracker (during MRI, Eyelink 1000 Plus Long-Range) focused on the right eye at a 1000 Hz sampling rate (except for 2 participants at 500 Hz). In both settings, the eye tracker was calibrated just prior to task onset for each person.

### Saliva collection and genotyping

Each participant provided a single saliva sample using DNA Genotek Oragene OGR-600 kits, at least 30 min after last consuming anything other than water. Genomic DNA was extracted with the MagMAX DNA Multi-Sample Ultra 2.0 Kit. APOE was genotyped by allele-specific real-time PCR (56), and the ADRA2B Δ301-303 insertion/deletion polymorphism by targeted amplicon next-generation sequencing; full protocols are in the **Supplement**. APOE and ADRA2B Del301-303 were the only loci genotyped (original hypotheses in F32 AG058479).

### MRI

We used a 3T GE Discovery MR750 with a 32-channel head coil at the Cornell MRI Facility. Ear plugs and headphones were used for ear protection. Anatomical data were acquired with a T1-weighted MPRAGE sequence (TR = 2600 ms; TE = 2.85 ms; TI = 1000 ms; 7° flip angle; phase acceleration factor 2; 1.0 mm isotropic voxels, 200 slices). Multi-echo echo planar imaging (EPI) sequences were used during the task (TR = 2500 ms; TEs = 14, 27, and 40 ms; multiband factor 2; phase acceleration factor 3; 80° flip angle; 1.83 x 1.83 x 2 mm voxels; 78 slices). 240 volumes were acquired (600 s), and the first 2 volumes were discarded leaving 238 volumes (595 s) for analysis.

### Pupil data preprocessing and quality

Pupil data were cleaned as in our previous work (55,57), following Kret & Sjak-Shie (58). For taVNS sessions, all missing samples were removed with a ±50 ms margin, samples with |Δpupil| > median(|Δ|) + 8×MAD were removed with a ±10 ms margin, pupil size was converted to mm, and gaps ≤ 500 ms were linearly interpolated. A trial was usable if no sample from 50 ms before to 2,400 ms after stimulus onset was missing after interpolation. Each usable trial was baseline-corrected to the mean of the 50 ms before onset, and the area under the curve (AUC) from 0 to 2.4 s was averaged across usable trials of each type and expressed as % of the participant’s dynamic range × s, using the dynamic range (1st to 99th percentile of cleaned pupil size) of the sham session for both sessions. Participants with fewer than 10 usable oddball trials in either session were excluded (n = 2, **Table 1**). Retained participants had a median of 30 usable oddball trials per session (range 19–30), which did not differ by carrier status (p = .31). For pupil-BOLD coupling analyses, the cleaned pupil trace was averaged in 50 ms bins aligned to the first retained fMRI volume and z-scored, and its first derivative was the pupil change regressor.

### fMRI data preprocessing and quality

Anatomical images were processed with FreeSurfer recon-all (59) and AFNI’s SSWarper2 (60), and multi-echo EPI was processed with AFNI (61), including multi-echo combination and denoising with tedana (62) and scaling to percent signal change. For cortical analysis, motion parameters were not entered as regressors; motion-related variance was removed by tedana, and volumes with > 5% of brain voxel outliers were censored. The Schaefer-200 atlas (63) was warped to each participant’s native space, and the mean time series of each parcel was extracted. Pupil size-BOLD coupling (“*size coupling”*) in each parcel was the Pearson correlation between the parcel time series and the pupil size regressor averaged within each TR with pupil leading BOLD by 2.25 s. Pupil change-BOLD coupling (“*change coupling”*) used the pupil change regressor with pupil leading by 5.0 s, the lags of peak pupil-BOLD cross-correlation in our previous work (55). Carriers and non-carriers did not differ in mean framewise motion (p = .96), number of censored volumes (p = .46) or mean absolute coupling across parcels (p = .72; Table 3).

**Table 3.** NET alignment of pupil change-BOLD coupling: specificity, robustness and group comparability.

| Row | Check | Model | Deletion carrier effect [95% CI] | P for deletion carrier effect |
|---|---|---|---|---|
| — | Primary result | NET alignment ~ carrier + age | b = 0.181 [0.048, 0.314] | **.010; adjusted Cohen's d = 1.21** |
| **Specificity** | Other neurotransmitter maps | Alignment with each of 18 other PET maps ~ carrier + age | all \|b\| ≤ 0.132, \|t\| ≤ 1.63 | all ≥ .117 |
|  | Pupil size-BOLD coupling | NET alignment of pupil size coupling ~ carrier + age | b = 0.026 [−0.127, 0.180] | .725 |
| **Robustness** | Highest 1, 2, 3 carriers removed | Primary model | b = 0.162 / 0.147 / 0.133 | **.018 / .032 / .052** |
|  | Highest-motion 1, 2, 3 participants removed | Primary model; motion = mean framewise motion (AFNI enorm) | b = 0.156 / 0.135 / 0.131 | **.030 / .051 / .062** |
|  | Leave-one-out | Primary model, each participant removed in turn (25 fits) | b = 0.156 to 0.219 | **.003 to .030 (25 of 25 < .05)** |
|  | Rank-based statistics | Mann-Whitney U = 26 | rank-biserial r = 0.62 | **.013** |
|  | Rank-based alignment | Spearman ρ (coupling map vs NET) ~ carrier + age | b = 0.179 [0.046, 0.311] | **.010** |
|  | Permutation | Primary model, carrier labels shuffled 10,000 times within age tertiles | b = 0.181 (observed) | **.013** |
|  | Spin test on the carrier difference | Carrier t recomputed against 5,000 spatially rotated NET maps | b = 0.181 (observed), t = 2.82 | **.010** |
|  | All recorded covariates at once | Primary model + sex + race + mean motion + censored volumes | b = 0.166 [0.008, 0.324] | **.041; adjusted Cohen's d = 1.11** |
|  | Age group instead of linear age | NET alignment ~ carrier + age group | b = 0.178 [0.044, 0.313] | **.012** |
| **Group comparability** | Head motion | Mean framewise motion ~ carrier + age | b = −0.002 mm | .96 |
|  | Censored volumes | Number censored ~ carrier + age | b = 2.2 volumes | .46 |
|  | Overall coupling strength | Mean \|r\| of coupling map ~ carrier + age | b = 0.008 | .72 |

### PET tracer analysis methods

Neurotransmitter receptor/transporter maps were drawn from Hansen et al., 2022 (64) using the neuromaps python package (65) as described previously (55). For each tracer, PET density was averaged within each of the 200 cortical parcels of the Schaefer 200 atlas, and each participant’s alignment with that tracer was the Pearson correlation across parcels between PET density and the participant’s pupil-BOLD coupling map. For spin tests, Alexander-Bloch (66) tests were used.

## Results

### Effects of transcutaneous vagus nerve stimulation on task-evoked pupillary responses in ADRA2B deletion carriers vs. non-carriers

Each person completed verum and sham taVNS sessions while performing an oddball task. To begin, we fitted an omnibus linear mixed effects model on task-evoked pupillary response area under the curve (AUC; one value per participant × treatment condition × trial type). It included fixed effects of treatment condition (verum vs. sham), trial type (oddball and bright, vs. standard), ADRA2B Δ301–303 deletion carrier status (carrier or non-carrier), and all their interactions, together with age, treatment order, their interactions with treatment condition, random intercepts for participant, and its interaction with trial type. We found significant effects of oddball trial type (b = 15.31, t(126) = 5.37, p < .001), bright trial type (b = −8.62, t(126) = −3.02, p = .003), and a three-way interaction between oddball trials, carrier status, and verum stimulation (b = 9.14, t(115) = 2.55, p = .012). This suggested that deletion carrier status specifically affected oddball responses during taVNS. Our primary outcome was therefore the within-participant Verum - Sham difference in oddball AUC. This metric captures the change in pupillary responses induced by verum taVNS.

We found that participants heterozygous or homozygous for the ADRA2B Δ301–303 deletion (hereafter “carriers”) responded differently to taVNS than non-carriers (b = 9.28 % dynamic range × s, 95% CI [3.08, 15.49], t(37) = 3.03, p = .004; N = 41, 16 non-carriers, 25 carriers, **Fig. 1A**). Carriers experienced more variable and positive responses to taVNS; non-carriers experienced a less variable and more negative response. The Verum - Sham difference also became more negative with age (b = −0.19 per year, [−0.34, −0.03], t(37) = −2.38, p = .023). Age was included as a covariate because of its known effect on the taVNS response in an overlapping cohort (36); its effect is consistent with that earlier report but is not an independent test of it. The carrier effect did not depend on age (carrier × age b = −0.10, t(36) = −0.66, p = .52; age slopes −0.18 in non-carriers and −0.22 in carriers). Carrier status and age each explained independent variance (unique R^2^ = .15 and .09 respectively).

The carrier difference was specific to oddball trials and to verum stimulation (**Table 2**). Carriers and non-carriers did not differ in the effect of taVNS on standard or bright trials (**Fig. 1B–C**), or in the sham session alone, but did differ in the verum session alone (**Fig. 1D**). The effect was robust to removing the strongest carrier responses and noisiest participants, leave-one-out analysis (41 of 41 significant), rank-based and permutation statistics, a different interpolation limit, adding sex, race, stimulation current difference and washout length as covariates, and adjusting for each participant’s sham response (**Table 2**). Carriers and non-carriers did not differ in baseline pupil diameter, stimulation current, washout duration or treatment order (**Table 2**).

Pupil traces demonstrated that carriers on average had larger oddball-evoked pupillary responses under verum taVNS, while non-carriers on average had smaller ones (**Fig 2F-H)**. In an exploratory cluster-based permutation test of the verum-sham trace (0-2400 ms, carrier + age + treatment order at each timepoint), a cluster of significant differences between carriers and non-carriers ran from 610 to 2390 ms after stimulus onset

**Fig. 2.**
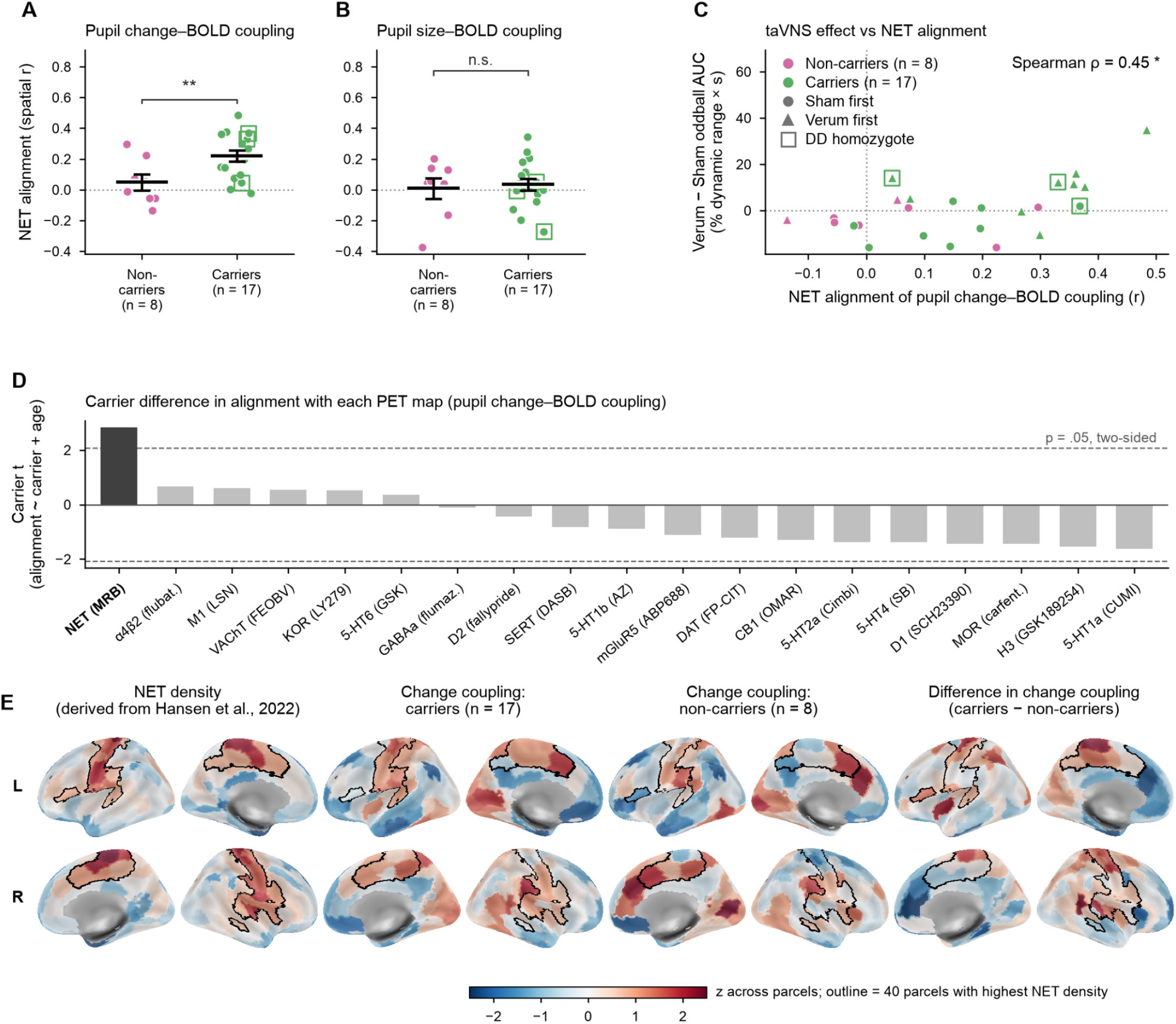
NET spatial alignment of pupil–BOLD coupling by ADRA2B carrier status. (A, B) Per-participant spatial correlation between NET density and pupil change-BOLD coupling (A) or pupil size-BOLD coupling (B) (N = 25; mean ±1 SEM shown; brackets show the carrier significance from alignment ∼ carrier + age). (C) NET spatial alignment of pupil change-BOLD coupling vs. the taVNS effect (Verum − Sham oddball AUC, % dynamic range × s). (D) Carrier t for alignment with each of 19 PET maps (alignment ∼ carrier + age); dashed lines, p = .05. (E) NET density and group-mean *change coupling* (correlation between pupil derivative and BOLD) in carriers, non-carriers and their difference, each z-scored across Schaefer-200 parcels; outline, the parcels with top 20% highest NET density. ^*^ p < .05, ^**^ p < .01.

The Verum - Sham difference was larger in participants who received Verum first (b = 8.28, [2.19, 14.37], t(37) = 2.76, p = .009) and larger responses were observed during the first session than the second. To investigate whether this effect was responsible for, or in addition to, the effect of carrier status, we added a carrier × order interaction. This showed that the treatment order effect differed by genotype and was only significant for carriers: interaction b = 19.81 [9.12, 30.50], p < .001; within carriers b = 15.81, p < .001; within non-carriers b = −3.80, p = .12 (**Fig. 1E**). In the 20 participants who did the verum taVNS condition first, 77% of carriers vs. 14% of non-carriers showed an increase in oddball AUC. In the 21 who did the sham condition first, 50% of carriers vs. 22% of non-carriers showed an increase. The carrier difference in the stimulation effect, averaged over the two treatment orders (i.e. sessions modeled as -0.5 and 0.5), was 9.93 [4.58, 15.28], t(36) = 3.77, p < .001, the same size as in the primary model. The carrier difference in the Verum response is therefore not an artifact of order, and the order effect is a second carrier-specific feature.

As a control, we tested whether APOE ε4 carrier status, added to the primary model, predicted the Verum − Sham difference: the per participant change in oddball-evoked pupillary responses induced by verum taVNS. APOE ε4 genotype had no significant effect (p = .57), and its inclusion did not decrease the effect of the ADRA2B deletion carrier status nor interact with it (for full model results see **Supplement**).

### BOLD correlates of pupillary responses in ADRA2B deletion carriers vs. non-carriers

We next conducted an exploratory analysis of the meaning of carriers’ and non-carriers’ pupillary activity on BOLD activity. Twenty-five of the taVNS participants (8 non-carriers, 17 carriers) also performed a version of the oddball task in the MRI scanner with pupillometry but without taVNS. We investigated BOLD coupling with continuous regressors for pupil size and change in pupil size, as well as task-evoked pupillary responses.

Oddball-evoked pupillary responses in the scanner did not differ between carriers and non-carriers (linear model on oddball AUC with fixed effects of carrier status and age group: t(21) = 1.54, p = .14; carrier × age group: F(2, 19) = 1.54, p = .24). Age group was used because oddball-evoked pupillary responses differ non-monotonically with age (36,55).

Two pupil-BOLD coupling maps were produced for each person: pupil size-BOLD coupling, and pupil derivative-BOLD coupling, hereafter “*size coupling*” and “*change coupling*” respectively, resulting in two parcellated correlation maps per person. We computed the parcelwise correlation coefficient between average NET density taken from existing PET maps (64,65), and each individual’s distinct coupling maps. Then, we constructed a linear model on each correlation coefficient, with fixed effects of carrier status and age, to determine whether carriers and non-carriers differed in how closely their coupling maps matched NET density. We found a significant effect of carrier status in how strongly change coupling correlated with NET density (b = 0.181, 95% CI [0.048, 0.314], t(22) = 2.82, p = .010, partial η^2^ = .27; N = 25, 8 non-carriers, 17 carriers, **Fig. 2A**): change coupling was more spatially NET-aligned in carriers than non-carriers (mean r with NET: carriers 0.220 (SD 0.150), non-carriers 0.048 (SD 0.148); greater than zero in carriers, p < .0001, but not in non-carriers, p = .39). By contrast, there was no effect of carrier status in how strongly size coupling spatially aligned with NET density (b = 0.026, 95% CI [−0.127, 0.180], t(22) = 0.36, p = .72, **Fig. 2B**), and it did not correlate significantly with NET density in either group.

Carriers and non-carriers differed only in the degree to which their change coupling correlated with NET density, not 18 other PET maps tested in follow-up (**Fig. 2D**). The NET alignment result held in every leave-one-out analysis (25 of 25 significant), with rank-based, permutation and spin-test statistics, and with sex, race, framewise motion and censored volumes as covariates. It remained significant after removing the one or two most NET-aligned carriers or the highest-motion participant, and fell to p = .051–.062 after removing three carriers or the two or three highest-motion participants (**Table 3**). Carriers and non-carriers did not differ in head motion, censored volumes, and overall pupil-BOLD coupling strength (as opposed to its spatial distribution, **Table 3**).

As a control, we included APOE ε4 carrier status with ADRA2B deletion carrier status, in the model predicting NET spatial alignment of change coupling. APOE ε4 carrier status was not a significant predictor of NET spatial alignment (p = .14), its presence in the model did not decrease the effect size of ADRA2B deletion carrier status (b = 0.187 with, 0.181 without), and the two did not interact (p = .92; for full results, see **Supplement)**.

In these 25 participants, we asked whether their individual response to taVNS (i.e., Verum - Sham difference in oddball-evoked AUC) could be predicted by the strength of their *change coupling* with NET density. We found that those with greater taVNS responsiveness also had greater NET spatial alignment of *change coupling* (Spearman ρ = .45, p = .023, **Fig. 2C**). This result was specific to carriers (within carriers ρ = .54, p = .026; within non-carriers ρ = .19, p = .65), and to the verum session (verum session alone ρ = .48, p = .015, sham session alone ρ = −.04, p = .83).

### Additional analyses

We investigated whether deletion carriers differed in their LC structural integrity, LC function (BOLD responses), cognitive performance, or heart rate variability. There were no significant effects found (see **Supplement**).

## Discussion

We identified a noradrenergic genetic variant, ADRA2B Δ301-303, that predicts significant variance in the effect of noninvasive VNS on task-evoked pupillary responses, which are used as a proxy for LC-NA activity. We also show, in those who completed fMRI, that the variant affects the degree to which pupil-linked cortical activity is concentrated in NA-innervated areas. Deletion carriers exhibited more variable responses to taVNS and were more likely to show increased pupillary responses than non-carriers. The effect of the deletion variant was specific to task-relevant oddball trials, rather than the pupillary light reflex, and to verum stimulation rather than sham. The putative mechanism of taVNS, as well other forms of VNS, has generally been described as noradrenergic (4,21–23,29–31,67–70); these results lend support to that hypothesis, and provide evidence that individual differences in genotype may be a critical contributor to efficacy of VNS (71).

Following De Quervain et al. (47), showing that carriers had enhanced emotional memory, studies from our laboratory extended these findings to emotional memory and perception (49,50), with convergent findings in an independent sample (52). These effects have been interpreted as potentiated noradrenergic modulation of attentional and perceptual processing (72,73). In our current study, the effect of the deletion variant on pupillary responses, as an index of task-evoked attentional orienting and arousal, was detectable only when comparing responses across conditions with putative differences in noradrenergic state, relative to baseline and a pupillary light reflex control. Deletion carriers showed stronger spatial correspondence with pupil-coupled neural activity and cortical NET expression, despite comparable pupillary responses across carriers and non-carriers without taVNS. These findings, along with previous evidence, suggest that the consequences of the deletion variant are salience-dependent, and concern the downstream consequences of NA release (e.g., action at NA receptors) in the brain. This suggests the ADRA2B variation may play a role in regulating moment-to-moment fluctuations of arousal and attention.

We found that ADRA2B deletion carriers are strongly affected by condition order, with larger pupillary responses on average in the first session compared to the second. This clearly points to the need, in future studies, to separate sessions on different days. However, the identification of this effect as being specific to deletion carriers again suggests arousal and attentional salience differences between genetic groups, uncovered by what may be a first-session novelty effect. It is not consistent with a carryover response, which would have increased sham responses in the verum-first group.

Due to our small sample size, we were unable to test for deletion dosage effects. We used a single-blind design and our active sham, while standard, has come into question in terms of whether it is truly an appropriate sham (74). Moreover, the precise mechanism for these genotype-dependent differences remains unknown. Although the deletion variant has been shown to reduce receptor desensitization *in vitro* (44), it has not been directly shown that this molecular change leads to increased action of NA in humans.

Predictors of any VNS response have been elusive, even in clinical contexts, and outcomes of taVNS heterogeneous. We previously found that taVNS resulted in reduced pupillary responses in older adults; here, including genotype, we find that carrier status explained more variance than age, with the two effects independent. Since increased pupillary responses likely reflect a gain in noradrenergic transmission, which is hypothesized to be neuroprotective (75), carrier status could predict which older adults may benefit from taVNS. Further, given that the use of VNS for epilepsy (22), migraine (27), stroke rehabilitation (69), and depression (30) is hypothesized to depend on NA modulation, it might be reasonable to check whether this common genetic variant, or other NA variants, affects VNS responsiveness in other contexts. Although pupillary responses are not a clinical readout and the relationship between a predictor of pupillary responses and a predictor of clinical efficacy is unknown, at least one recent paper has proposed pupillary responses as a biomarker of VNS responsiveness to epilepsy treatment (76).

## Contributions

CRediT authorship contribution statement: **E. Riley:** Conceptualization, Methodology, Software, Validation, Formal analysis, Investigation, Resources, Data curation, Writing - original draft, Writing - review & editing, Visualization, Supervision, Project administration, Funding acquisition **S. A. Mabry:** Methodology, Software, Formal analysis, Investigation **M. MacMillan:** Methodology, Investigation **L. Ko:** Project Administration, Writing - original draft **F. Middleton:** Methodology, Investigation, Resources, Writing - original draft, Writing – review & editing **E. De Rosa:** Resources, Supervision, Funding acquisition, Writing – review & editing **A. Anderson:** Conceptualization, Resources, Supervision, Funding acquisition, Writing – review & editing

## Data and code availability

Data and code are available on request

## Use of artificial intelligence tools

The initial data analysis code was written by the corresponding author between 2022 and 2026. Beginning in 2026, Anthropic models (Claude; models Opus 5.0/5.5, Fable 5.1) were utilized to optimize code. The corresponding author used the same models to verify the accuracy of each number reported in the text against the analysis outputs and to populate data tables and model results directly from analysis outputs. AI tools were not used to write or edit the manuscript. The authors take full responsibility for the integrity and content of this work.

