## Supplement for "ADRA2B genetic polymorphism modulates vagus nerve stimulation response"

#### Contents:

1. Supplementary results
    a. Effect of ADRA2B deletion on oddball task performance
    b. Effect of ADRA2B deletion and APOE4 ε4 variants on locus coeruleus structural integrity
    c. Effect of ADRA2B deletion and APOE4 ε4 variants on LC BOLD
    d. Effect of ADRA2B deletion and APOE4 ε4 variants on cognitive performance
    e. Effect of ADRA2B deletion and APOE4 ε4 variants on heart rate variability (HRV)
    f. Detailed results of APOE4 ε4 negative controls presented in main text
2. Methods
    a. Participant info for additional analyses
    b. Genetic analysis
    c. LC-sensitive MRI
    d. LC BOLD
    e. Physiological measurement for calculation of heart rate variability

#### Supplementary Results

##### Effect of ADRA2B deletion on oddball task performance

We investigated the effects of deletion carrier status on task performance (linear models with fixed effects of carrier status, age and treatment order; N = 39; two participants did not have behavioral response files). Under sham stimulation, oddball detection was at ceiling (99.5% of targets detected) and false alarms were rare (0.34% of standard trials), and neither these nor median reaction time differed between carriers and non-carriers (detection $p = .71$, false alarms $p = .32$, reaction time $p = .37$). Carrier status did not predict the Verum − Sham change in any measure (detection $p = .89$, false alarms $p = .34$, reaction time $p = .95$).

##### Effect of ADRA2B deletion and APOE4 ε4 variants on locus coeruleus structural integrity

We investigated whether deletion carriers vs. non-carriers differed in their locus coeruleus structural integrity, cognitive performance, or heart rate variability. Using LC-sensitive MRI, we found no difference between groups in LC intensity (maximum LC intensity relative to pons, linear model with fixed effects of carrier status and age group: $b = -0.088$, 95% CI [−0.196, 0.020], $t(125) = -1.62$, $p = .11$; carrier × age group: $F(2, 123) = 0.31$, $p = .73$; N = 129, 58 non-carriers, 71 carriers). Because the TSE sequence changed after a scanner software upgrade (n = 38 old sequence, 91 new sequence), we repeated the model with sequence as a covariate; the carrier effect was unchanged ($b = -0.088$, $p = .11$; carrier × sequence $p = .95$).

APOE ε4 carrier status was also not associated with LC intensity (same model with ε4 carrier status in place of deletion carrier status: $b = -0.001$, 95% CI [−0.113, 0.111], $t(125) = -0.02$, $p = .98$; ε4 × age group: $F(2, 123) = 1.09$, $p = .34$; N = 129, 80 ε4 non-carriers, 49 ε4 carriers).

#### Effect of ADRA2B deletion and APOE4 ε4 variants on LC BOLD

We also measured LC BOLD oddball responses (FIR at 2.5 s after stimulus onset, which we have previously shown is the time of peak response for this task (1)), with full physiological denoising, within individually specific LC ROIs, and found no difference between carriers and non-carriers ($b = -0.059$ % signal, 95% CI [−0.213, 0.095], $t(22) = -0.80$, $p = .43$; N = 25, 8 non-carriers, 17 carriers).

LC BOLD oddball responses also did not differ by APOE ε4 carrier status ($b = 0.043$ % signal, 95% CI [−0.118, 0.204], $t(22) = 0.55$, $p = .59$; N = 25, 17 ε4 non-carriers, 8 ε4 carriers).

#### Effect of ADRA2B deletion and APOE4 ε4 variants on cognitive performance

On the NIH Toolbox Cognition Battery (N = 186, 89 non-carriers, 97 carriers), we found no significant difference between groups, after correction for multiple comparisons, although deletion carriers had higher scores on 4 out of 5 subtests used. Scores that were higher in carriers included Dimensional Change Card Sort, Pattern Comparison Processing Speed, Flanker Inhibitory Control, and List Sorting Working Memory. Picture Sequence Memory scores were lower in carriers. For List Sorting Working Memory, carriers had significantly higher scores before correction for multiple comparisons but not after ($p = .020$, means 101.7 vs 107.0; Holm-corrected $p = .098$, allele dose for List Sorting: $p = .038$, but not monotonic [II 101.7, ID 107.2, DD 106.2; n = 88 / 82 / 15]).

APOE ε4 carriers (N = 186, 126 ε4 non-carriers, 60 ε4 carriers; N = 185 for List Sorting) did not differ from non-carriers on any NIH Toolbox subtest before or after correction for multiple comparisons (linear models with fixed effects of ε4 carrier status and age; all $p \geq .10$, all Holm-corrected $p \geq .51$). ε4 carriers had lower scores on Dimensional Change Card Sort, Picture Sequence Memory and Flanker Inhibitory Control, and higher scores on Pattern Comparison Processing Speed and List Sorting Working Memory. The largest difference was in Dimensional Change Card Sort (means 110.6 vs 105.5; $b = -5.12$, $t(183) = -1.65$, $p = .10$, Holm-corrected $p = .51$).

#### Effect of ADRA2B deletion and APOE4 ε4 variants on heart rate variability (HRV)

Using heart rate data collected in the MRI scanner, we measured heart rate variability (ln RMSSD) and found no difference ($b = 0.061$, 95% CI [−0.590, 0.711], $t(20) = 0.19$, $p = .85$, N = 23, 8 non-carriers, 15 carriers).

HRV also did not differ by APOE ε4 carrier status (ln RMSSD: $b = 0.509$, 95% CI [−0.158, 1.176], $t(20) = 1.59$, $p = .13$, N = 23, 16 ε4 non-carriers, 7 ε4 carriers).

#### Detailed results of APOE4 ε4 negative controls presented in main text

We tested whether inclusion of APOE ε4 carrier status (i.e., the presence of at least one copy of APOE ε4, vs. no copies of ε4), along with ADRA2B deletion carrier status, predicted our primary measure: the per participant change in oddball-evoked pupillary responses induced by verum taVNS. It had no significant effect (APOE ε4 $b = 1.92$ % dynamic range × s, $t(36) = 0.57$, $p = .57$; N = 41, 29 without ε4, 12 with ε4), and its inclusion did not decrease the effect of the ADRA2B deletion carrier status (ADRA2B $b = 9.28$, $t(37) = 3.03$, $p = .004$ without APOE; $b = 9.18$, $t(36) = 2.96$, $p = .005$ with APOE). It also did not significantly interact with ADRA2B deletion carrier status in predicting the per participant effect of taVNS (ADRA2B × APOE ε4 $b = 1.73$, $p = .82$).

We tested whether APOE ε4 carrier status, along with ADRA2B deletion carrier status, was a significant predictor of NET alignment of change coupling, including age as a covariate as before. APOE ε4 carrier status was not a significant predictor of NET alignment ($b = -0.099$, $t(21) = -1.52$, $p = .14$; N = 25, 17 without ε4, 8 with ε4), and its presence in the model did not decrease the size of the effect of ADRA2B deletion carrier status (ADRA2B $b = 0.181$, $t(22) = 2.82$, $p = .010$ without APOE; $b = 0.187$, $t(21) = 3.00$, $p = .007$ with APOE). The two did not interact (ADRA2B × APOE ε4 $b = 0.015$, $p = .92$; carrier − non-carrier difference in NET alignment 0.18 among both ε4 non-carriers and ε4 carriers). We additionally tested whether APOE ε4 carriers and noncarriers differed in their alignment of size coupling or change coupling to any of 18 other PET maps. No map differed by ε4 carrier status at $p < .05$ for either change coupling (smallest $p = .085$, VAChT) or size coupling (smallest $p = .056$, NET); all Holm-corrected $p = 1$.

### Methods

#### Participants

**Table S1. Participant information for additional analyses**

| Cohort | Final sample size | Original sample size | Age: M (SD), range | Sex | Race and Ethnicity | Dates of data collection | Cause of exclusions |
|---|---|---|---|---|---|---|---|
| **NIH Toolbox Cognition Toolbox + genotype** | 186 (89 non-carriers, 97 carriers) | 186 | 51.7 (18.7), range 19-86 | 129 F, 57 M | W 123, A 22, B 35, Multi 4, AI/AN 2; H 7 | Feb 2019 - Dec 2025 | none |
| **LC-sensitive structural MRI + genotype** | 129 (58 non-carriers, 71 carriers) | 131 | 54.2 (17.0), range 19-82 | 98 F, 31 M | W 88, A 11, B 25, Multi 3, AI/AN 2; H 6 | Mar 2019 - Dec 2025 | 2 excluded after visual QC (artifacts) |

| | | | | | | | |
|---|---|---|---|---|---|---|---|
| **Heart rate recorded during fMRI + genotype** | 23 (8 non-carriers, 15 carriers) | 25 | 46.4 (17.7), range 19-78 | 17 F, 6 M | W 15, A 6, B 2; H 0 | Jun 2023 - Sep 2025 | 1 no physiological data; 1 excluded after visual QC (artifacts) |

#### Genetic analysis

##### *DNA Extraction*

Expectorated samples were stored at room temperature, sealed and protected from light, for 1-12 months before being moved to 4°C. For processing samples were mixed thoroughly by inversion and incubated at 50°C for a minimum of 1 hour in their original collection tubes to complete cell lysis and digest nucleoprotein complexes. Automated genomic DNA isolation was performed from 200 µL aliquots of the stabilized saliva using the MagMAX DNA Multi-Sample Ultra 2.0 Kit (Applied Biosystems / Thermo Fisher Scientific) on a KingFisher Flex Purification System according to the manufacturer protocol. Briefly, samples were treated with Proteinase K and Lysis Solution, followed by the addition of magnetic Binding Beads and isopropanol/binding enhancer to facilitate DNA capture. The KingFisher Flex automated the sequential binding, magnetic collection, and transfer of the bead-bound DNA through two distinct wash solutions and an 80% ethanol wash to eliminate residual proteins, salts, and PCR inhibitors. Purified genomic DNA was eluted into 50 µL of Elution Solution, checked for purity and concentration using a Nanodrop, diluted to 10 ng per µL with PCR grade nuclease-free water and stored at −20°C until further analysis.

##### *Apolipoprotein E (APOE) Genotyping*

Genotyping was performed using allele-specific real-time PCR adapted from the method described by Zhong et al. (2016), incorporating a dual-quenched fluorogenic probe design. All primers and probes were synthesized by Integrated DNA Technologies (Coralville, IA). Allelic discrimination utilized the published forward (F) and reverse (R) allele-specific primer sets (e2 F/R: GCGGACATGGAGGACGTGT / CCTGGTACACTGCCAGGCA; e3 F/R: CGGACATGGAGGACGTGT / CTGGTACACTGCCAGGCG; e4 F/R: CGGACATGGAGGACGTGC / CTGGTACACTGCCAGGCG) alongside a modified common internal probe incorporating an internal ZEN quencher and 3' Iowa Black quencher (5′-FAM/CAGCTCCTC /ZEN/ GGTGCTCTGGC/3IAbRQSp-3′). Three separate PCR reactions were performed for each DNA sample, interrogating the possible presence of 1 specific ApoE allele per reaction. PCR amplifications were assembled in 20 µL total reaction volumes containing 20 ng of purified genomic DNA, TaqMan Environmental Master Mix 2.0 (Applied Biosystems), 1 µL each of forward and reverse primers from 10 µM stock solutions (final concentration 500 nM each), and 1 µL of the fluorogenic probe from a 5 µM stock solution. Amplification and real-time detection were conducted on a CFX Opus Real-Time PCR System (Bio-Rad Laboratories) using the following cycling profile: initial hot-start and enzyme activation at 95°C for 10 min, followed by 42 cycles of denaturation at 95°C for 20s, annealing at 59°C for 20s, and extension with fluorescence acquisition at 62°C for 20s. Genotypes (APOE ε2, ε3, and ε4 alleles) were assigned by evaluating differential cycle quantification (Cq) values across the allele-specific reaction wells using Bio-Rad CFX Maestro software.

##### *ADRA2B Genotyping*

Due to complexities surrounding the methodologies and reporting of sequence variations in the ADRA2B gene, we performed genotypic analysis of a functional insertion/deletion polymorphism within the ADRA2B gene (spanning the polymorphic glutamic acid repeat region) using targeted amplicon next-generation sequencing (NGS). A 330-bp target region on human reference genome GRCh38/hg38 (chr2:96,115,082–96,115,411) was amplified using custom primers (Forward: 5′-CCACTGCCCACCTATAGCAC-3′; Reverse: 5′-GTGGACTTCGAGTGTCCGTT-3′). PCR reactions were assembled in 20 µL volumes containing Q5 Hot Start High-Fidelity 2X Master Mix (New England Biolabs) and 20 nanograms purified genomic DNA. Following amplification, amplicons were purified using MagMAX PureBind magnetic beads on a KingFisher Flex system and eluted in 50 µL of PCR grade nuclease-free Tris-HCl (pH 8.5). Multiplexed sequencing libraries were prepared from 40 ng of purified amplicon DNA using the ExpressPlex Library Preparation Kit (seqWell) according to the manufacturer's instructions. Resulting libraries were examined on an Agilent TapeStation. Paired-end sequencing (2 × 150 bp) was carried out on an Illumina NextSeq 2000 platform using a P1 reagent kit. Demultiplexed FASTQ reads were aligned to the GRCh38 human reference genome using BWA-MEM, and local realignment along with insertion/deletion variant calling and genotyping was performed using the Genome Analysis Toolkit (GATK) HaplotypeCaller algorithm.

#### LC-sensitive MRI

To measure LC structural integrity, we used a 2D T1-weighted fast spin-echo sequence (ETL = 3, TR = 600 ms, TE = 14 ms, flip angle = 90°, field-of-view 22 x 22 cm, phase FOV = 80%, 6 averages, 0.43 x 0.43 x 3mm resolution with the acquisition plane perpendicular to the brainstem). Following our previously published methods for processing this data (2,3), the LC was located within each scan. Briefly, two trained raters traced the LC following the algorithm described in (3), and voxels included in both tracings were included in the final ROI. If the Dice coefficient was not higher than 0.6, both raters redid the tracing.

#### LC BOLD

BOLD responses within SegmentAAN LC labels were estimated, in native space, for each trial type with a finite impulse response (TENT) model with knots every 2.5 s from −2.5 to 15 s after stimulus onset. Polynomial drift terms and nuisance regressors for cardiac and respiratory noise modelled from the physiological recordings (RETROICOR cardiac and respiratory phase regressors (4), heart rate convolved with the cardiac response function, and respiration volume per time convolved with the respiration response function), together with the first five principal components of the 4th ventricle signal (5), were added with outlier volumes censored as in the whole-brain analysis.

Bilateral LC signals were partial-volume weighted and averaged across hemispheres before model fitting.

#### Physiological measurement for calculation of heart rate variability

During fMRI, heart rate and respiratory activity were continuously recorded at 100 Hz (cardiac) and 25 Hz (respiratory). Heart rate was obtained from a photoplethysmography (PPG) sensor placed on the nondominant index finger, and respiration was recorded with a pneumatic respiration belt. Heart rate variability was computed from the PPG recording of the oddball run; the first 5 s were discarded and the following 595 s analysed, matching the retained fMRI volumes. Systolic peaks were detected with NeuroKit2 (6). Inter-beat intervals 1.5-2.5 times the median length were split in two (missed beats), intervals shorter than 0.65 times the median were merged with the following interval (false peaks), and remaining intervals deviating more than 25% from the median or lying outside 0.3-2.0 s were replaced by cubic-spline interpolation. RMSSD was computed on the corrected series and natural-log transformed for analysis because of skewness. Raw and corrected interval series were inspected visually for each participant, and participants whose PPG could not be corrected reliably were excluded (**Table S1**).
